# Vestigial like 4 regulates the adipogenesis of classical brown adipose tissue

**DOI:** 10.1101/2024.07.09.602788

**Authors:** Pingzhu Zhou, Chase W. Kessinger, Fei Gu, Amanda Davenport, Justin S. King, Genyu Wang, Steven G. Negron, Bart Deplancke, William T. Pu, Zhiqiang Lin

## Abstract

Brown adipose tissue (BAT) is mammals’ primary non-shivering thermogenesis organ, and the molecular mechanisms regulating BAT growth and adipogenesis are largely unknown. The Hippo-YAP pathway has been well-known for controlling organ size, and Vestigial like 4 (VGLL4) is a transcriptional regulator that modulates the Hippo-YAP pathway by competing against YAP for binding to TEAD proteins. In this study, we dissected the function of VGLL4 in regulating BAT development. We generated a conventional *Vgll4* mutant mouse line, in which the two Tondu (TDU) domains of VGLL4 were disrupted. We found that deletion of the TDU domains of VGLL4 resulted in perinatal lethality and paucity of the interscapular BAT. Histological and magnetic resonance imaging studies confirmed that the adipogenesis of BAT was impaired in *Vgll4* mutants. Adeno-associated virus (AAV) mediated, brown adipocyte-specific overexpression of VGLL4 increased BAT volume and protected the adult male mice from acute cold stress. Genomic studies suggest that VGLL4/TEAD1 complex directly regulates the myogenic and adipogenic gene expression programs of BAT. In conclusion, our data identify VGLL4 as a previously unrecognized adipogenesis factor that regulates classical BAT development.

## Introduction

Obesity is a global epidemic that contributes to a number of chronic disease conditions, such as type 2 diabetes mellitus, non-alcoholic fatty liver, cardiovascular disease, and cancers^1^. In mammals, the adipose tissue comprises white and brown adipose tissue (WAT and BAT, respectively) ^2^. WAT stores triglycerides in adipocytes, and its expansion is a hallmark of obesity. On the other hand, BAT burns triglycerides and glucose to generate heat ^3^. Based on its developmental origin, BAT is further divided into classical BAT and white adipocyte-derived brown adipocytes (beige/brite cells) ^4^. In humans, BAT is abundant in infants, and decreases with age ^5^. Recently, the discovery of functional BAT and beige cells in adults ^6^ raises the possibility of treating obesity by activating BAT or increasing the browning of WAT ^7^. Thus understanding the molecular mechanisms that control BAT growth and function is critical for developing therapeutic strategies to treat obesity.

Both white adipocytes and classical brown adipocytes (BACs) are derived from mesenchymal stem cells ^8^. BAC adipogenesis generally includes four steps. First, mesenchymal stem cells differentiate into myogenic factor 5 (Myf5)-expressing progenitor cells. Next, Myf5^+^ progenitor cells differentiate into brown preadipocytes, which then differentiate into UCP1-positive naive BACs. Finally, naive BACs develop into lipid-loaded and mitochondria abundant mature BACs ^9^. In mice, the interscapular BAT depot appears at embryonic day 15.5 (E15.5), and expands rapidly during fetal and neonatal development. BAT lipogenesis is minimal in murine embryos and increases remarkably after birth ^10,11^.

The genetic programs controlling BAT adipogenesis are well documented. Transcription factors, including peroxisome proliferator-activated receptor gamma (PPARγ), CCAAT/ enhancer-binding proteins (specifically CEBPβ/δ), and PRD1-BF-1-RIZ1 homologous domain containing protein-16 (PRDM16), are key regulators of BAC differentiation ^8,9,12^. Conditional deletion of *Pparg* in adipocytes reduced both WAT and BAT ^13^. Mice missing both CEBPβ and CEBPδ had no lipid accumulation in BAT, and primary embryonic fibroblasts lacking both of these factors failed to undergo adipocyte differentiation ^14^. PRDM16 is a crucial regulator of myogenic and brown adipogenic lineage differentiation, because ectopic expression of PRDM16 in myoblasts activated the BAC differentiation program, which relied on the interaction between PRDM16 and CEBPβ/δ ^15^.

The Hippo-YAP kinase signaling pathway is well known for controlling organ growth ^16^. In mammals, upstream signaling pathways activate the Hippo kinase cascade, comprised of MST1/2, LATS1/2, and the scaffold protein Salvador (Sav). Activation of this kinase cascade results in phosphorylation and inactivation of YAP and WWTR1 (more commonly referred to as TAZ), orthologous transcriptional co-activators that are terminal transcriptional effectors of this pathway. YAP/TAZ interact with TEAD family transcription factors to regulate downstream target gene expression, including genes that control cell proliferation ^17^. Several studies have implicated Hippo-YAP signaling in WAT adipogenesis. In the 3T3-L1 preadipocyte cell line, which has been widely used to study WAT adipocyte differentiation ^18^, overexpression or knock-down of Hippo kinases activated or suppressed adipogenesis, respectively ^19,20^, while knock-down of YAP/TAZ or TEAD4 augmented WAT adipogenesis ^21,22^. YAP/TAZ-TEAD complex also plays an important role in regulating BAT adipogenesis. BAT-specific YAP/TAZ double heterozygous mice (YTU) had reduced BAT size and impaired metabolic activity at 4 weeks of age. At 20 weeks of age, compared with the BAT of the age matched control mice, the YTU BAT was two times larger, which was due to excessive lipid accumulation in the BACs ^23^. However, the underlying molecular mechanism of how YAP/TAZ-TEAD complex regulates BAT adipogenesis has not been well documented.

In addition to inhibition by the Hippo kinase cascade, the activity of the YAP/TAZ-TEAD complex is also suppressed by Vestigial like 4 (VGLL4), a transcriptional regulator that competes with YAP/TAZ for binding to TEAD ^24,25^. In 3T3-L1 cells, loss of either TEAD4 or VGLL4 increased adipogenesis; overexpressing TEAD4 suppressed adipogenesis, which was abolished by knocking down VGLL4 ^22^. These data highlight the importance of VGLL4 as a crucial regulator of adipogenesis in vitro and raise the question whether VGLL4 regulates adipogenesis in vivo.

Previously, we generated a *Vgll4* mutant mouse allele, *Vgll4^d^*^46^, in which a 46 bp deletion caused a frameshift mutation and disrupted the TDU domain that mediates TEAD binding. *Vgll4^d46/d46^* (*Vko*) mice could not prevail and died shortly after birth ^26^. Using interscapular BAT as a classical BAT example, here we show that *Vko* mice had much less BAT than their littermate controls, and that VGLL4 is essential for the normal BAT adipogenesis gene expression program. Additionally, overexpressing VGLL4 in postnatal BAT increased BAT volume and protected the mice from acute cold stress. In summary, our study reveals that VGLL4 is a crucial adipogenesis factor that controls BAT growth and function, thus advancing the knowledge of BAT development.

## Results

### Mutating VGLL4 causes BAT paucity

We have previously reported that whole-body depletion of VGLL4 led to perinatal death ^26^. While characterizing the phenotype of the *Vgll4* knockout (*Vko*) pups, we noticed that the *Vko* newborn pups had consistent dents in the interscapular regions, which led us to examine the interscapular BAT depot. Impressively, the *Vko* pups had much smaller BAT depots than their littermate controls (Fig. 1A). By performing qRT-PCR with the P0 BAT we confirmed the depletion of *Vgll4* (Suppl. Fig. 1A).

**Figure 1.**
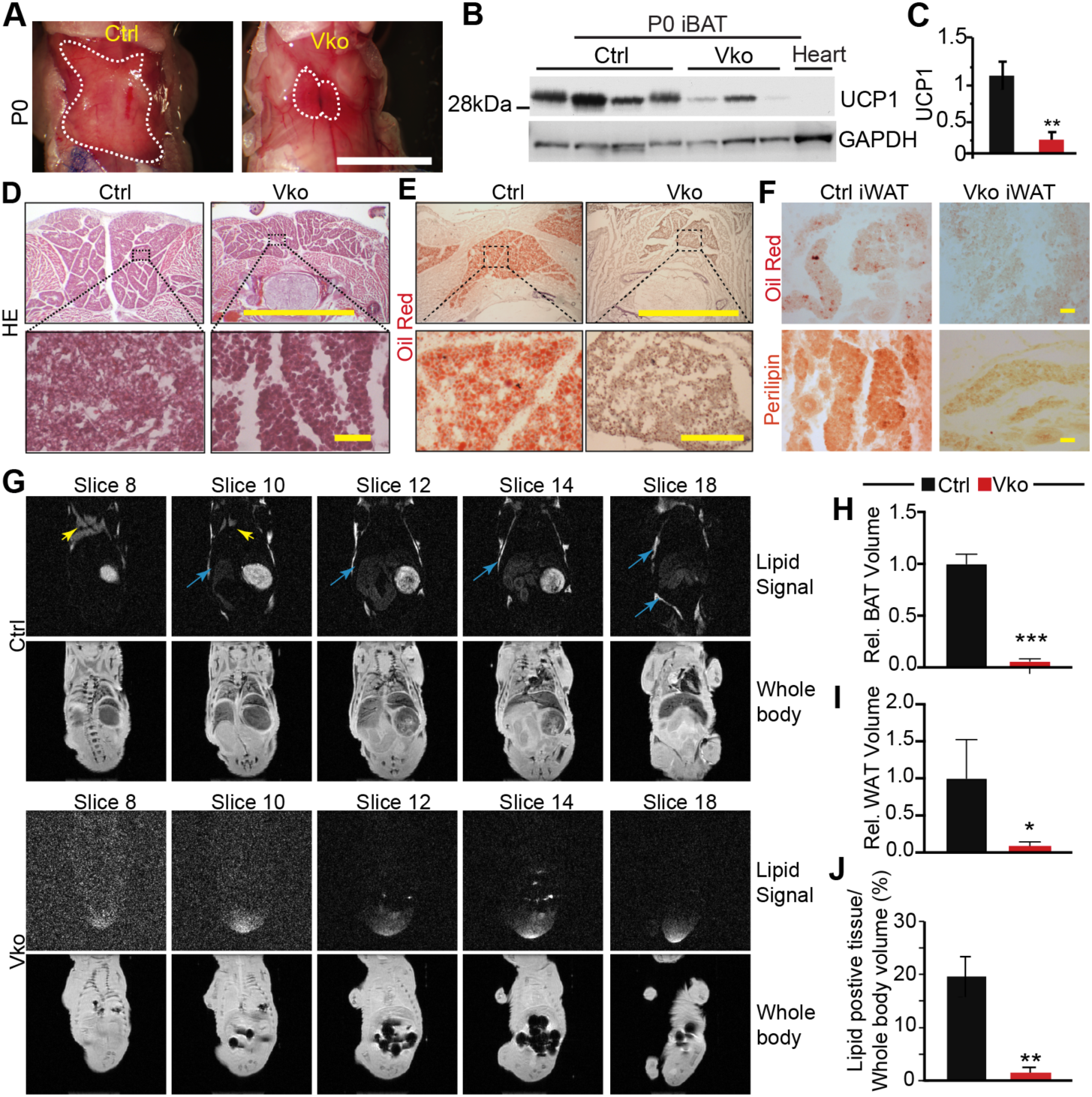
Classical brown adipose tissue growth requires VGLL4. **A.** Gross morphology of classical interscapular brown adipose tissue at P0. Dotted lines indicate the BAT depot. Scale bar=4 mm. **B**. UCP1 immunoblot with P0 BAT protein. Heart protein was used as negative control. **C**. Densitometry quantification of UCP1. Protein levels were normalized to GAPDH. N=4. Student’s t test, **, P<0.01. **D**. HE staining of BAT. Top panel, scale bar =4 mm; bottom panel, scale bar=50 µm. **E.** Oil Red O staining of BAT. Top panel, bar= 4 mm; bottom panel, scale bar=1 mm. **F.** Oil Red O staining and Perilipin immunohistological staining of iWAT. Scale bar = 50 µm **G**. Visualization of whole body trunk for lipid content by a series of MRI images using P0 mouse pups. Water signal was suppressed in the lipid only images. Blue and yellow arrows indicate BAT and and subcutaneous WAT, respectively. **H.** BAT lipid content comparison. **I**. WAT lipid content comparison. **J**. Quantification of the total lipid content. The percentage was calculated by normalizing the lipid positive tissue volume to the body trunk volume. Student t-test: *P<0.05, n=4.

To elucidate the cellular mechanism of BAT paucity, we first examined the BAC size and found reduced BAC size in *Vko* compared with littermate controls (Suppl. Fig. 1B and 1C). We then measured the proliferation rate of the UCP1-positive BACs with EdU, a thymidine analogue that labels proliferating cells. *Vko* BACs had a comparable EdU incorporation rate to controls (Suppl. Fig. 2D and 2E). Consistent with the malformation of BAT, the expression of BAT marker UCP1 was reduced in the *Vko* mutants (Fig. 1B and 1C), suggesting that loss of VGLL4 may either interfere with preadipocyte-to-BAC differentiation or impair BAC maturation.

To determine if the BAT abnormalities preceded birth, we collected embryos at E18.5 and examined the interscapular BAT depot. Compared with wild type controls, the *Vgll4* heterozygous (*Vhet*) embryos had no obvious growth defects, as reflected by the normal body weight (Suppl. Fig. 1F). The *Vko* mutants were smaller than both wild type and *Vhet* embryos; however, the weight of the major internal organs, such as the heart and lungs, normalized to body weight, was not affected (Suppl. Fig. 1G and 1H). Because we observed no phenotypic difference between *Vhet* and wild type mice, we used both of these genotypes as controls. In the control embryos, the interscapular regions were filled with BAT covered by white adipocytes (Suppl. Fig. 1I); however, the *Vko* embryos had much smaller interscapular BAT depots (Suppl. Fig. 1I), demonstrating that BAT paucity occurs in *Vko* during embryonic development.

We further examined the BAT histology and lipid deposition in the neonatal *Vko* mutants. The BAT depot was thinner in *Vko* than in controls (Fig. 1D, top row). The nuclei were more dense in *Vko* BAT, and *Vko* BACs had less cytoplasmic vacuoles (Fig. 1D, bottom row).

Oil Red O staining revealed that lipid droplets were abundant in control BAT but rarely detected in the *Vko* BAT (Fig. 1E), suggesting that VGLL4 is essential for BAT lipogenesis.

Similar to BAT, the inguinal subcutaneous white adipose tissue (iWAT) contained minimal lipid droplets in *Vko*, whereas lipid droplets were readily detected in control iWAT (Fig. 1F). Perilipin is a protein enriched on the surface of lipid droplets in differentiated adipocytes ^27^. Compared with control iWAT, *Vko* iWAT had much less perilipin immunoreactivity (Fig. 1F). To more broadly assess the effect of VGLL4 mutation on lipid deposition, we measured the lipid composition of the chest and abdomen of control and *Vko* pups with magnetic resonance imaging (MRI). Lipid signals were detected in the interscapular and subcutaneous regions of control but not *Vko* pups (Fig. 1G). Quantification of the relative volume of BAT and subcutaneous WAT revealed that both these adipose depots were much smaller in the *Vko* pups (Fig. 1H and 1I). Additionally, quantification of the lipid signals contained in the chest and abdomen tissues demonstrated a significant decrease of lipid content in the *Vko* mutants (Fig. 1J, *Vko* 1.49% vs Ctrl 19.59%, p=0.01). Together, these data suggest that VGLL4 regulates adipogenesis in both BAT and subcutaneous WAT.

### Loss of VGLL4 disrupts the adipogenesis gene expression program of BAT

To determine whether VGLL4 inactivation changed the BAT gene expression program, we performed RNA sequencing with BAT collected from P0 pups. Compared with littermate controls, 1217 and 978 genes were significantly down and up-regulated in the *Vko* BAT, respectively (P_adj_<0.05 and absolute FC>1.5; Fig. 2A, Dataset S1). Whole genome Gene Set Enrichment Analysis (GESA) analysis ^28^ showed that myogenesis, oxidative phosphorylation, fatty acid metabolism, and adipogenesis were the top four biological processes enriched in the wild type BAT (Fig. 2B; Suppl. Fig.2A). Out of 191 genes in the adipogenesis gene set, 51 were down-regulated in *Vko* (Fig. 2C). We used qRT-PCR to validate the expression of *Acsl1, Adipor2, Cidea* and *Dgat1,* and confirmed that all these genes except *Adipor2* were decreased in *Vko* BAT (Fig. 2D).

**Figure 2.**
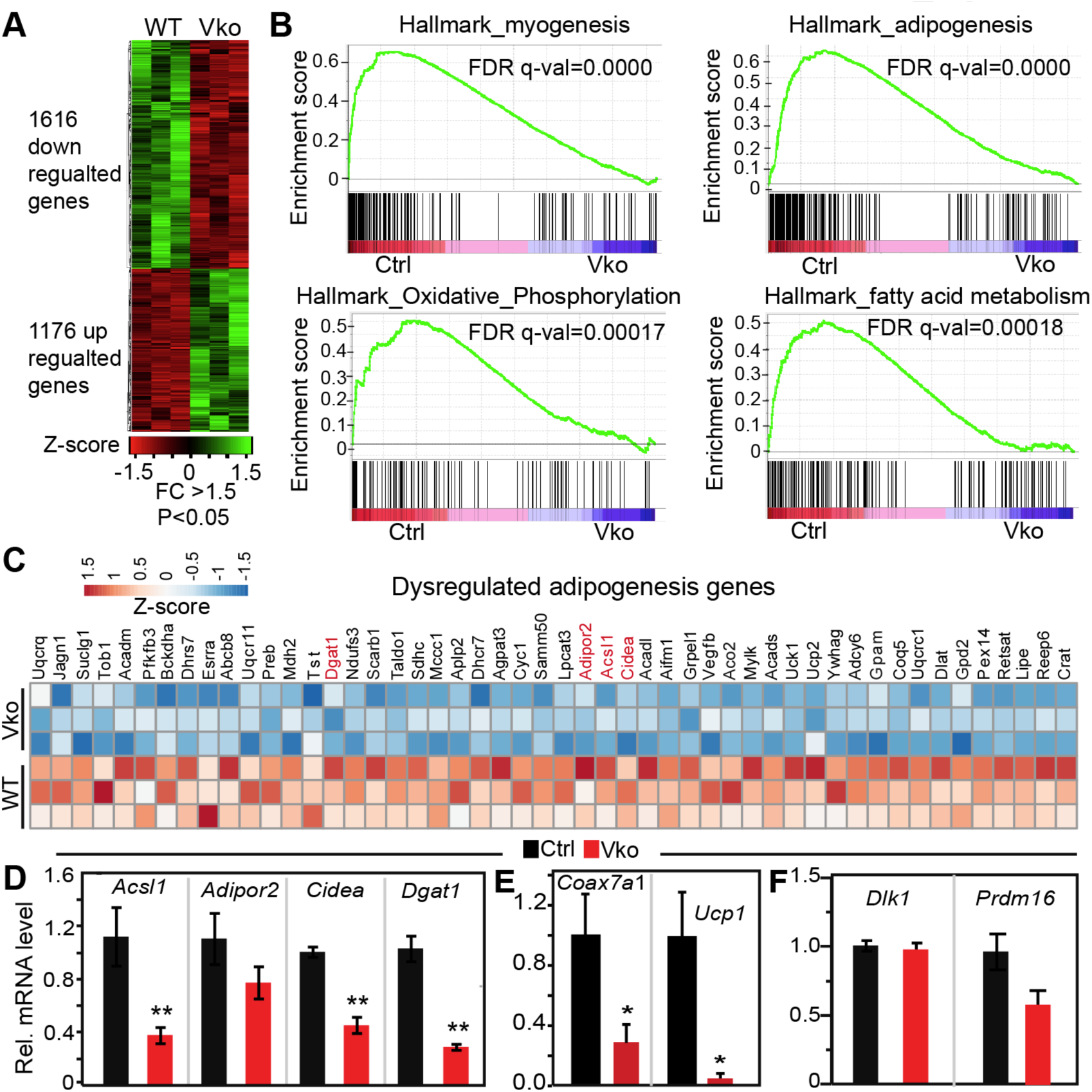
Loss of VGLL4 disrupts the adipogenesis gene expression program of BAT. **A.** Heat map of differentially expressed genes (DEG). RNA sequencing data from P0 BAT was analyzed with Deseq. N=3. **B**. Gene set enrichment analysis (GESA). GESA was carried out with the Deseq2-derived gene expression data set. The top four gene sets enriched in wild-type BAT were shown. **C**. A heat map of differentially expressed fatty acid metabolism/ adipogenesis genes. **D**. qRT-PCR validation of selected lipogenesis genes dysregulated in *Vko* BAT. **E-F.** qRT-PCR measurement of brown adipocyte differentiation-related genes. D, E and F, N=5. Student t-test, *, P<0.05. All the tested mRNAs were normalized by endogenous 36B4 mRNA.

To assess the BAC differentiation stage that is regulated by VGLL4, we took advantage of published gene expression profiles of BACs at different stages of differentiation ^29^. In this study, 960 protein coding genes were found to be differentially expressed during pre- adipocyte to BAC differentiation. Among these 960 genes, 856 genes were detected in our RNA-seq data. Based on the expression data of these 856 genes, we performed hierarchical clustering to determine which stage of differentiation was most similar to P0 control or *Vko* BAT. Both the control and *Vko* BAT gene expression data clustered with that of the differentiated BACs (Suppl. Fig. 2B), suggestive a minor role of VGLL4 during brown preadipocyte to BAC differentiation.

We further measured the expression of two BAC-enriched genes (*Ucp1* and *Cox7a1*) and two BAC differentiation-related genes (*Dlk1/Pref-1* and *Prdm16*). In comparison with control BAT, *Vko* BAT had lower expression of *Ucp1* and *Cox7a1* (Fig. 2E) and similar expression of *Dlk1* and *Prdm16* (Fig. 2F).

### Deletion of VGLL4 activates IGF-Akt pathway

VGLL4 is a transcriptional regulator that blunts the interaction between YAP and TEAD proteins ^25^. It has been reported that mice doubly heterozygous for YAP and its paralog TAZ had disrupted BAT growth, and that YAP/TAZ coordinated with TEAD1 to drive UCP1 expression in BACs ^23^. These published data suggest that TEAD1 is involved in the development of BAT. To test the correlation between TEAD1 expression and BAT development, we examined *Tead1* mRNA and TEAD1 protein levels in the BAT at different developmental stages. *Tead1* mRNA and TEAD1 protein were both high in postnatal day 1 (P1) BAT and then quickly decreased to a very low level at P4 (Fig. 3A; Suppl. Fig. 3A). Different from *Tead1*, *Vgll4* mRNA level was decreased from P1 to P15 and then increased from P15 to P30 (Supp. Fig. 3B).

**Figure 3.**
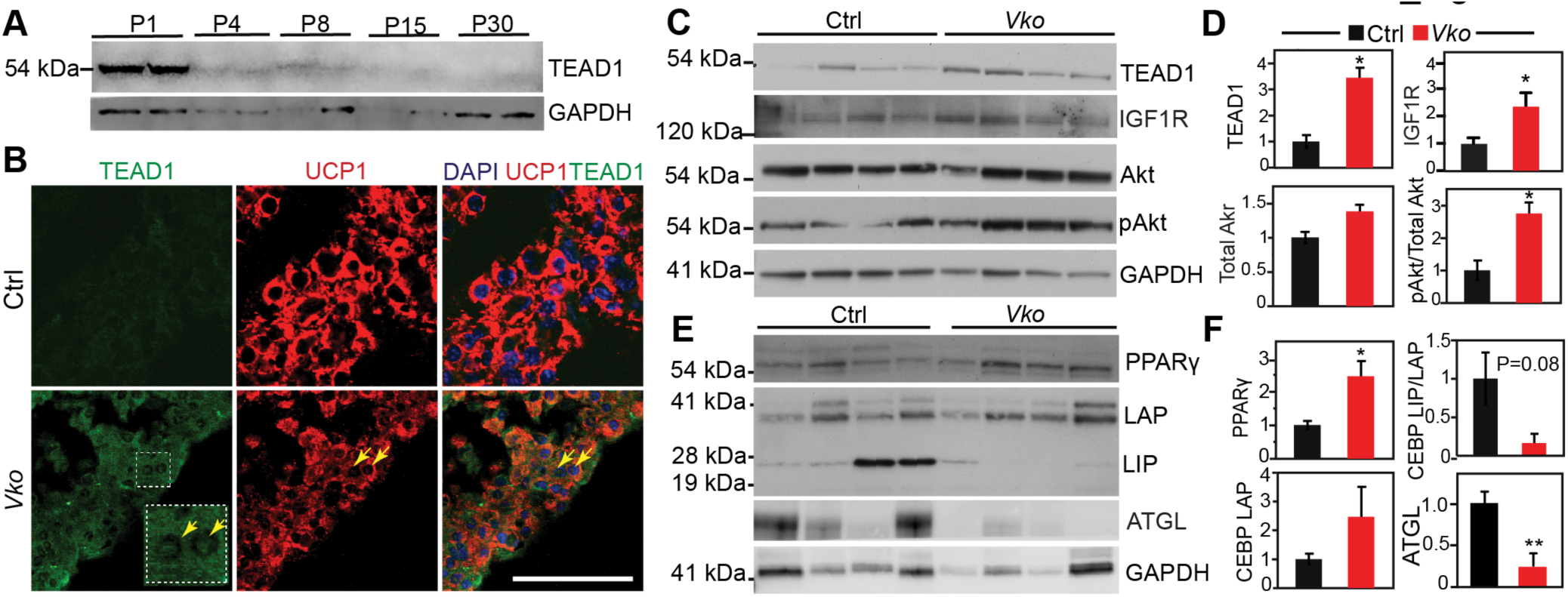
Loss of VGLL4 activates TEAD1. **A.** Western blot analysis of TEAD1 expression in BAT. BAT proteins collected from mice of different ages were compared by western blot. GAPDH was used as loading control. **B**. Immunofluorescence images of UCP1 and TEAD1 stained BAT sections. Yellow arrows indicate brown adipocytes enriched with nuclear TEAD1. Scale bar = 50 µm. **C**. Western blot analysis of TEAD1 and the crucial IGF1-Akt pathway components. **D**. Densitometry quantification of TEAD1, IGF1R, total Akt and phosphorylated Akt. **E**. Western blot analysis of crucial adipogenesis regulators. **F**. Densitometry quantification of PPARγ, C-LAP, C-LIP and ATGL. P0 interscapular BAT protein was used for western blots. D and F, protein levels were normalized to GAPDH. N=4. Student’s t test, *, P<0.05; **, P<0.01.

We previously showed that activation of VGLL4 decreased TEAD1 protein level in the heart ^25^ and knocking out VGLL4 increased TEAD1^26^. We then checked whether inactivation of VGLL4 affected TEAD1 protein expression in BAT. Immunofluorescence staining showed that TEAD1 expression and nuclear abundance was both increased in the BACs of neonatal *Vko* (Fig. 3B). In line with these in situ observations, western blots confirmed that *Vko* BAT had more TEAD1 than littermate controls (Fig. 3C and 3D). The YAP/TEAD complex regulates the expression of IGFR-Akt pathway components ^30,31^. Consistent with the upregulation of TEAD1, the *Vko* pups had increased IGF1R and higher phospho Akt/Akt ratio (Fig. 3C and 3D). These data suggest that VGLL4 ablation increases the transcriptional activity of the TEAD1/YAP complex in BAT.

Next, we measured the protein levels of PPAR-γ and CEBPβ, two transcriptional factors central to the regulation of adipogenesis ^32,33^. PPARγ was significantly increased in *Vko* BAT (Fig. 3E and 3F ), suggesting that loss of VGLL4 does not impair BAT adipogenesis by decreasing PPARγ expression. CEBPβ plays a central role in adipocyte differentiation ^33^, and it has several N-terminally truncated isoforms, such as liver-enriched activator protein (LAP) and liver-enriched inhibitory protein (LIP). CEBPβ-LAP interacts with PRDM16 to direct the differentiation of myoblastic precursors into BACs ^34^, and CEBPβ-LIP rescued the growth defects of *Cebpb* knockout mice ^35^, suggesting that CEBP-LIP and CEBP-LAP have overlapping functions. Compared to control BAT, CEBPβ-LAP expression in *Vko* BAT was similar, but CEBPβ-LIP was significantly lower (Fig. 3E and 3F).

Adipose triglyceride lipase (ATGL) is a well-known lipolysis enzyme and highly expressed in the adipose tissues of mice and humans ^36^. It recently was shown to regulate the synthesis of branched fatty acid esters of hydroxy fatty acids (FAHFAs) ^37^, a class of newly discovered lipids that have favorable metabolic and anti-inflammatory effects ^38^. In *Vko* BAT, ATGL expression was very low compared to control BAT (Fig. 3E and 3F), further suggesting that loss of VGLL4 impairs the development of BAT.

### VGLL4/TEAD1 complex regulates the expression of BAT myogenesis genes

We suspected that VGLL4 might interact with TEAD1 to regulate BAC gene expression. To test this hypothesis, we used a genomic approach to identify the genes that are regulated by both VGLL4 and TEAD1.

Because TEAD1 expression was high in the young newborn pups (Fig. 3A), we used P0 BAT to identify genes directly bound by TEAD1 using TEAD1 chromatin immunoprecipitation followed by next generation sequencing (ChIP seq). *Tead1^fb^* knockin mice that express FLAG and BIO epitope tagged TEAD1 ^25^ were crossed to BirA transgenic mice ^39^ to generate *Tead1^fb^*;*BirA* mice, in which the endogenous TEAD1^fb^ protein was biotylated by BirA.

TEAD1 bioChIP-seq of P0 *Tead1^fb^;BirA* BAT revealed 4234 peaks that mapped to 2834 genes, with 371 peaks localized in the promoter region (0 to -5kb upstream of the transcriptional starting site), and the rest of the peaks spreading out in the gene bodies and distal noncoding regions (Suppl. Fig. 4A). De novo motif discovery ^40^ identified the TEAD binding motif as the mostly enriched motif (Fig. 4A), thus confirming the reliability of our ChIP seq data set. The other top five transcriptional factors that were co-enriched in the TEAD1 ChIP peaks were Nuclear factor 1 (NF1), CCCTC-Binding Factor (CTCF), CEBP, Early B Cell Factor transcription factor 1 (EBF1) and Myocyte Enhancer Factor 2C (MEF2c) (Fig. 4A). Among these five transcriptional factors, both NF1 and CEBP protein families are essential for BAT adipogenesis ^14,41^, suggesting that TEAD1 may coordinate with these transcriptional factors to regulate BAT development.

**Figure 4.**
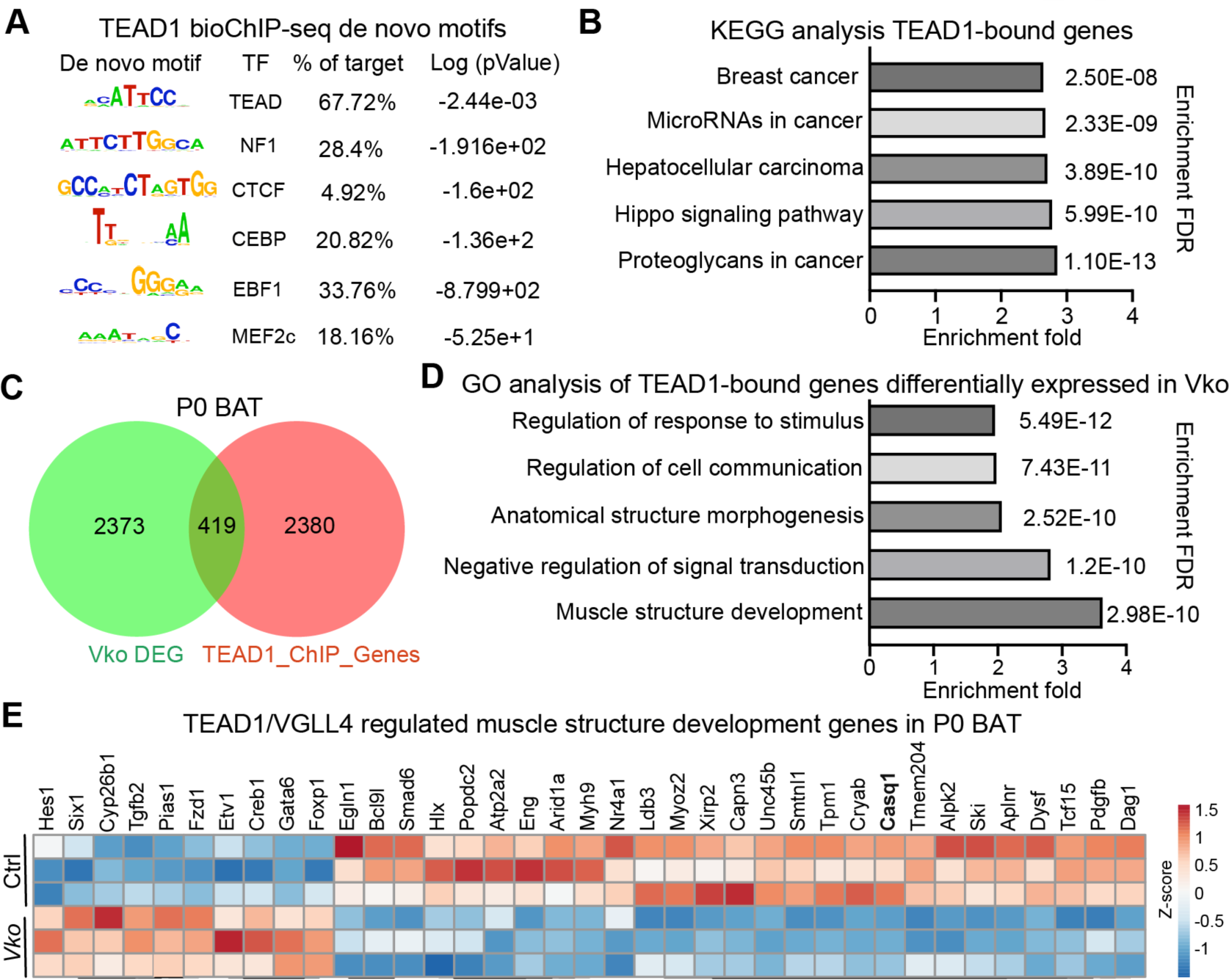
Identification of VGLL4/TEAD1 target genes in BAT. **A**. De novo motif discovery analysis within the identified TEAD1 binding sites. P0 BAT was used for TEAD1 ChIP seq. **B**. KEGG analysis of TEAD1 bound genes. The bar graph shows the top five enriched signaling pathways. **C**. Venn diagram showing the relationship between TEAD1 bound genes and differentially expressed genes in *Vko* (*Vko* DEG) brown adipose tissue. **D**. Gene Ontology analysis of the TEAD1/VGLL4 target genes. The 419 genes both dysregulated in *Vko* and bound by TEAD1 were applied to Shiny GO analysis. The top 5 enriched biological processes are listed in the bar graph. **E.** Heat map of differentially expressed VGLL4/TEAD1 regulated myogenic genes. RNA sequencing data from P0 BAT were presented. N=3.

Pathway analysis with the 2834 TEAD1-bound genes showed that the Hippo signaling pathway was among the top five enriched pathways (Fig. 4B). By comparing the *Vko* BAT differentially regulated genes (DEG) with the TEAD1-bound genes, we identified 419 overlapped genes that might be directly regulated by VGLL4 and TEAD1 (Fig. 4C). Gene ontology term analysis ^42^ of these 419 genes revealed that muscle structure development was the most enriched biological process (Fig. 4D). 37 TEAD1-bound genes were differentially expressed in *Vko* BAT and related to muscle structure development (Fig. 4E). Myogenic gene expression is a molecular signature that separates BACs from white adipocytes ^43^. These data suggest that the VGLL4/TEAD1 complex regulates the expression of myogenic genes in BAT.

In addition to regulating myogenic genes, TEAD1 also bound to crucial lipogenesis genes, such as *Fabp4, Lpl, Fabp3, Cd36, Prdm16, Cav1, Slc27a1,* and *Ppargc1b* (Suppl. Fig. 4B). qRT-PCR tests confirmed that *Cav1*, *Fabp3*, *Slc27a1,* and *Ppargc1b* were all down regulated in the *Vko* BAT (Suppl. Fig. 4C).

### VGLL4 alleviates TEAD1 suppression of CEBPβ transcriptional activity

Calsequestrin1 (*Casq1*) encodes a high-capacity calcium-binding protein that acts as a Ca^2+^ storage/buffer molecule in the sarcoplasmic reticulum of skeletal muscle cells ^44^. *Casq1* is also expressed in BAT, and down regulation of *Casq1* is associated with decreased BAT thermogenic capacity ^45^. Unbiased *Vko* BAT gene expression profiling and TEAD1 ChIP-seq analysis identified *Casq1* as a direct target of the VGLL4/TEAD1 complex (Fig. 4E). As shown in Fig. 5A, TEAD1 directly bound to the promoter region of *Casq1*. In BAT, *Vgll4* inactivation robustly decreased *Casq1* mRNA (Fig. 5B). TEAD1 ChIP-seq peak region encompassed 480 base pairs (bp) of the *Casq1* promoter. Analysis of mouse and human sequences using the evolutionary conserved regions (ECR) genome browser ^46^ showed that the sequence is highly conserved (Fig. 5C) and contains one CEBP binding motif and one TEAD1 binding motif, both of which were identical between mouse and human (Fig. 5D).

**Figure 5.**
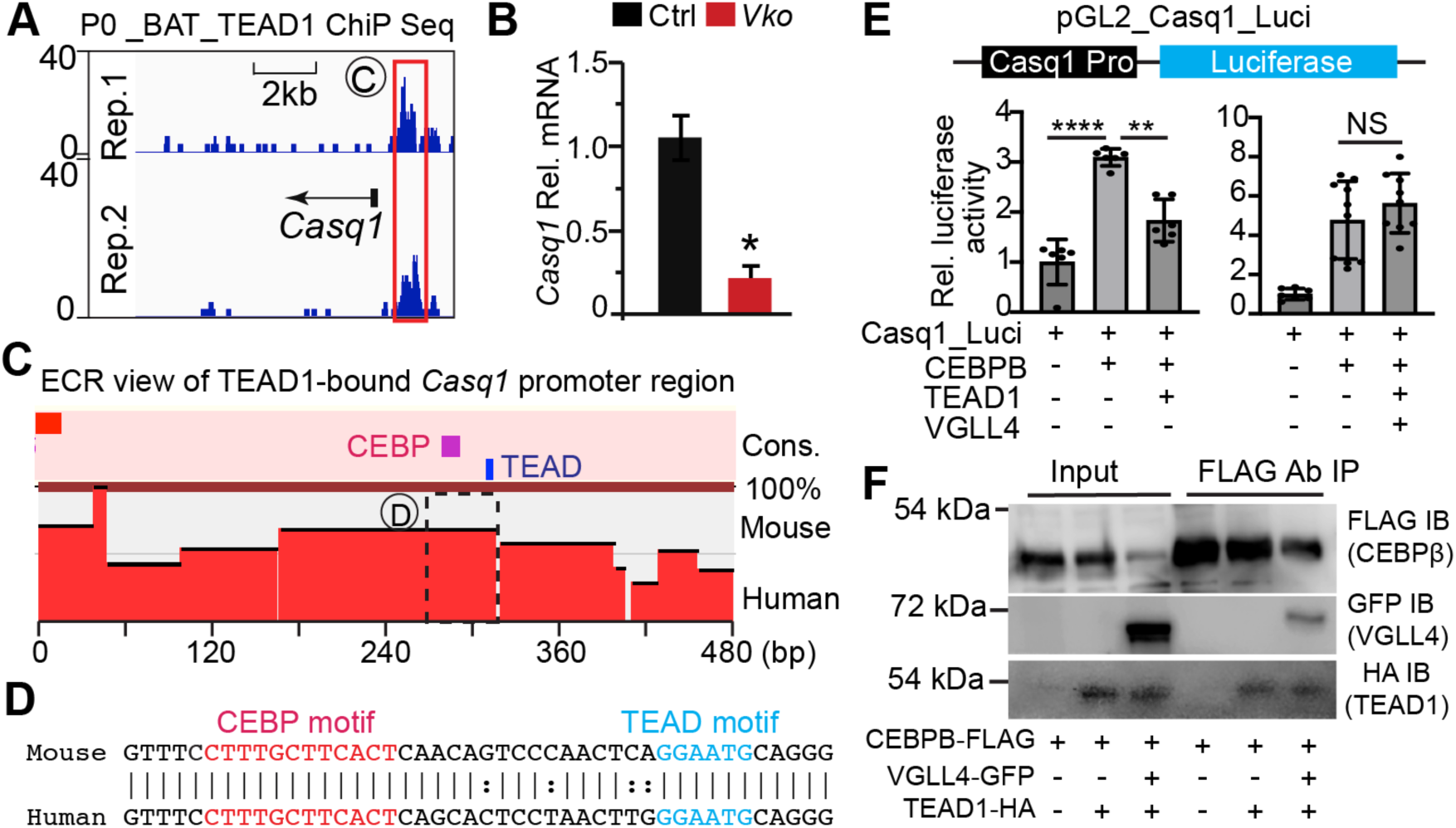
VGLL4 alleviates TEAD1 suppression of CEBP transcriptional activity. **A.** Genome browser view showing ChIP-seq of TEAD1 occupancy near the transcriptional start site (TSS) or *Casq1*. ChIP-seq data from two biological replicates are shown. **B.** qRT-PCR measurement of *Casq1*. P0 control and *Vko* BAT mRNA was used for qRT-PCR analysis. All the tested mRNAs were normalized by endogenous 36B4 mRNA. N=4. Student t-test, *, P<0.05. **C**. Evolutionary conservation of genomes (ECR) browser view of the TEAD1-bound *Casq1* promoter region. **D**. Alignment of the mouse and human *Casq1* promoter sequences enclosed by dotted rectangle in C. Conserved CEBP and TEAD binding motifs are highlighted. **E**. Luciferase reporter assay. The TEAD1-bound *Casq1* promoter sequence was cloned into pGL2 to generate Caqs1_Luci reporter plasmid. Indicated plasmids were transfected into HEK293T cells and *Casq1* promoter activity was measured by dual-luciferase assay. N=6. One-way ANOVA followed by multiple comparison analysis, ****, p<0.0001; ***, p<0.001. **F**. Co-immunoprecipitation (Co-IP) assay. Indicated plasmids were transfected into HEK293T cells. Flag antibody was used for Co-IP.

To study the activity of murine *Casq1* promoter, we constructed Casq1-Luci, in which the 480 bp TEAD1-bound region was positioned upstream of the luciferase gene in the promoterless pGL2 basic vector (Fig. 5E). In HEK293T cells, CEBPβ increased Casq1-Luci reporter activity, which was attenuated by further addition of TEAD1. VGLL4 did not affect CEBPβ transcriptional activity, but it abolished TEAD1 suppression of CEBPβ (Fig. 5E).

We performed co-immunoprecipitation (Co-IP) assays in HEK293T cells to characterize the relationships between CEBPβ, VGLL4, and TEAD1. Plasmids expressing TEAD1-HA, VGLL4-GFP and CEBPB-FLAG were co-transfected into HEK293T cells in different combinations. In the absence of TEAD1-HA, CEBPβ-FLAG pulled down VGLL4-GFP (Suppl. Fig.5); in the absence of VGLL4-GFP, CEBPβ-FLAG pulled down TEAD1-HA (Fig. 5F). When these three proteins were co-expressed in HEK293T cells, CEBPβ pulled down both VGLL4 and TEAD1 (Fig. 5F). These data suggest that TEAD1 and VGLL4 each bind to CEBPβ in a non-exclusive manner.

Altogether, these data suggest that TEAD1 suppresses CEBPβ activation of *Casq1* expression, and that VGLL4 favors *Casq1* expression by abolishing TEAD1 suppression of CEBPβ.

### Construction and validation of a BAC-specific AAV construct

AAV-mediated transduction of adipose tissue is a convenient tool for studying adipose pathophysiology, and AAV serotype 9 has a very high transduction efficiency for BAT ^47^. To facilitate gene-function studies in BAT, we generated a new AAV construct in which brown adipose tissue-specific cis-regulatory elements (BCE) derived from murine *Ucp1* drive cargo gene expression (Suppl. Fig. 6). To test the specificity and activity of BCE, we cloned either luciferase or GFP into this construct (Fig. 6A). 3-day-old pups (P3) were treated with AAV9.BCE.Luci and AAV9.BCE.GFP. 6 weeks post AAV delivery, we examined the tissue specificity of these AAV constructs (Fig. 6B). In vivo bioluminescence imaging showed that luciferase activity was limited to the interscapular region (Fig. 6C). Overlay of the luciferase signal onto whole animal microCT data confirmed that it originated from BAT (Fig. 6D). Histological sections of interscapular adipose tissue confirmed that BCE-driven GFP was expressed in BACs and not in the adjacent white adipocytes (Fig. 6E).

**Figure 6.**
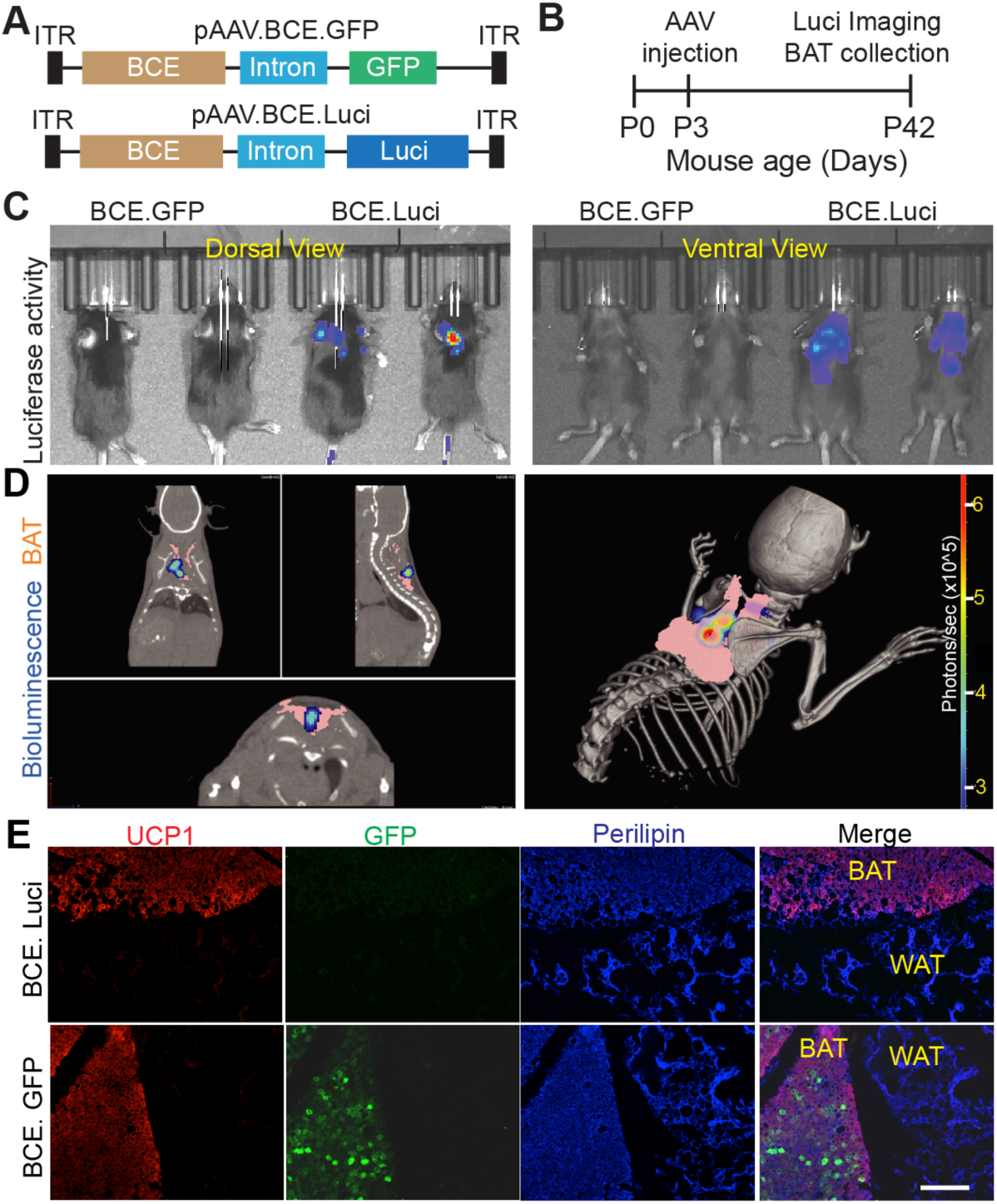
AAV9.BCE specifically expresses cargo genes in brown adipocytes. **A.** Schematic representation of pAAV.BCE.Luci and pAAV.BCE.GFP construct. Luciferase or GFP gene was placed downstream of BCE. One chimera intron of human beta-globin and immunoglobulin heavy chain genes was placed between BCE and cargo genes to increase cargo gene expression. ITR: inverted terminal repeat sequence. **B**. Experimental design. AAV9.BCE.luci or AAV9.BCE.GFP was injected into 3 days old pups (P3) subcutaneously. At 42 days after birth (P42), bioluminescence activity was measured with PerkinElmer in vivo imaging system (IVIS). **C**. Representative bioluminescence images generated by IVIS system. Both dorsal and ventral views were presented. **D**. Representative coregistration of bioluminescence signal and *microCT* segmented interscapular brown adipose tissue from an AAV.BCE.Luci mouse highlights the origin of the biolumenscence signal within the BAT volume. Pink: brown adipose tissue. Rainbow, bioluminescence signal positive tissues. **E**. Immunofluoresence images of BAT. BAT was collected at P42 and used for immunofluorescence staining. UCP1, BAT marker protein. Perilipin: adipose tissue marker (BAT and WAT) protein. Scale bar=200 µm

### Overexpressing VGLL4 in BACs promotes BAT growth

Because loss of VGLL4 caused BAT paucity (Fig. 1), we asked whether overexpressing VGLL4 in BACs promotes BAT growth.

To answer this question we generated AAV.BCE.VGLL4^GFP^ (BCE.VGLL4^GFP^), in which BAC-specific BCE regulatory elements drive the expression of GFP tagged human VGLL4. We treated p3 pups with BCE.VGLL4^GFP^ and assessed BAT volume at P42 using microCT (Fig. 7A). BCE.GFP was used as a control. Compared with BCE.GFP treated mice, BCE.VGLL4^GFP^ mice showed significantly higher BAT volume (102.20 ± 2.77 vs 84.16 ± 1.86 mm^3^, Fig. 7B), whereas the body weight of these two cohorts animals was not distinguishable (Fig. 7C). Consistent with the *microCT* measurements, the BAT weight to body weight ratio was significantly higher for BCE.VGLL4^GFP^ compared to BCE.GFP (Fig. 7D).

**Figure 7.**
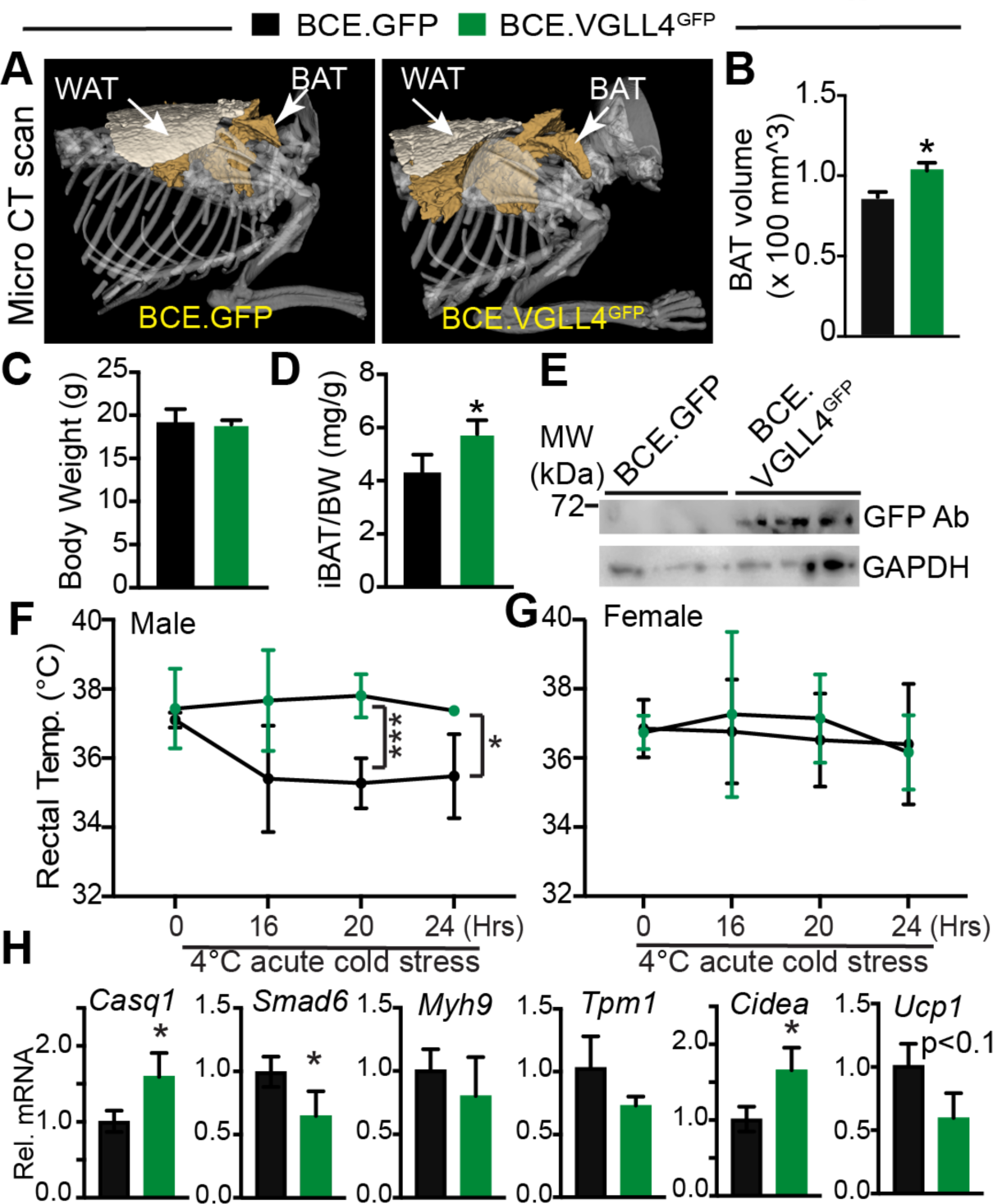
AAV9.BCE.VGLL4 promotes BAT growth. **A**. Representative volumetric images generated by microCT scan highlight the BAT (brown) and WAT (white) in the interscapular regions of the mice. Bones were labeled as grey. **B**. Quantification of BAT volume. **C.** Body weight. **D**. Interscapular BAT and body weight ratio. N=3. Student t-test, *, p<0.05. **E**. VGLL4 immunoblot with P42 BAT protein. GAPDH was used as internal control. **F-G.** Acute cold-stress test. BCE.GFP or BCE.VGLL4^GFP^ was injected into 3 days old pups (P3) subcutaneously. At 42 days after birth (P42), mice were treated with acute cold stress (4°C) for 24 hours. E, male mice rectal temperature. BCE.GFP, n=4; BCE.VGLL4^GFP^, n=3. Student t-test, ***, p<0.001; *, p<0.05. F, female mice core temperature, n=4 for each group. **H.** qRT-PCR measurement of gene expression. P42 BAT mRNA was used for qRT-PCR analysis. All the tested mRNAs were normalized by endogenous 36B4 mRNA. N=4. Student t-test, *, P<0.05.

We collected the BAT tissue to confirm the expression of VGLL4^GFP^. Western blot detected the expression of VGLL4^GFP^ in the BCE.VGLL4^GFP^ transduced mice (Fig. 7E), and in situ immunofluorescence staining showed that VGLL4^GFP^ signals were enriched in the nuclei of BACs (Suppl. Fig. 7A).

Because lipogenesis-driven BAC enlargement is one of the underlying mechanisms of postnatal BAT growth ^11^, we suspected that VGLL4 might promote BAT growth by increasing BAC size. We tested this hypothesis by staining BAT sections with Phalloidin, which labels the filamentous actin cytoskeleton ^48^. Compared to BCE.GFP, BCE.VGLL4^GFP^ significantly increased BAC size (Suppl. Fig. 7B and 7C), which was associated with the increase of lipid content as visualized by BODIPY ( a neutral lipid dye) staining (Suppl. Fig. 7D and 7E). To corroborate the in vivo results, we overexpressed VGLL4 in an immortalized brown preadipocyte (IBA) cell line ^29^ and tested VGLL4’s adiogenetic effects. Compared with GFP control, VGLL4 promoted lipid deposition in differentiated IBA cells (Suppl. Fig. 7F). Correspondingly, qRT-PCR results demonstrated the robust increase of three lipogenesis genes: *Fasn*, *Cidea* and *Acsl1* (Suppl. Fig. 7G). Thus, our in vitro and in vivo data strongly suggest that activation of VGLL4 promotes BAT growth by increasing BAC lipogenesis.

Thermogenesis is one of the biological functions of BAT. We asked whether activating VGLL4 in BAT affected whole body thermogenesis. To address this issue, we transduced P3 wild type mouse pups with BCE.VGLL4^GFP^ and performed acute cold stress test 6 weeks post AAV transduction. Age-matched BCE.GFP transduced mice were used as control. BCE.VGLL4^GFP^ and BCE.GFP mice were applied to acute 4°C cold stress for 24 hours. The core temperature of these two cohorts of mice were indistinguishable under regular housing temperature (∼22 °C). In the first 20-hour acute cold stress period, the core temperature of male BCE.GFP mice decreased from 37.10 ± 0.21 °C to 35.28 ± 0.72 °C; however, the core temperature of the male BCE.VGLL4^GFP^ mice did not change (37.43 ± 1.15 vs 37.80 ± 0.62) (Fig. 7F). In contrast to the male mice, the female mice did not show a core temperature difference between BCE.GFP and BCE.VGLL4^GFP^ groups (Fig. 7G).

On the molecular level, we collected BAT after acute cold stress and measured the expression of several VGLL4/TEAD1 target genes, including *Casq1, Myh9, Smad6,* and *Tpm1.* qRT-PCR results showed that VGLL4 increased *Casq1* and decreased *Smad6* (Fig. 7H). In line with our observation that VGLL4 promoted lipogenesis, *Cidea* expression was increased in BCE.VGLL4^GFP^ BAT. Although VGLL4 activation increased BAT thermogenesis, we found that VGLL4 decreased *Ucp1* expression, rather than increasing it (Fig. 7H).

## Discussion

In this study, we found that germ-line deletion of the VGLL4 TDU domains impaired BAT growth. Our data suggest that BAT adipogenesis requires VGLL4.

From fetus to early life, the development of BACs encompasses four stages: 1) mesenchymal stem cells differentiate into myogenic factor 5 (Myf5)-expressing progenitor cells; 2) Myf5^+^ progenitor cells differentiate into brown preadipocytes; 3) preadipocytes differentiate into UCP1-positive naive BACs; 4) naive BACs develop into lipid-loaded and mitochondria abundant mature BACs ^9^. Our current data advocate the hypothesis that VGLL4 regulates BAT growth by orchestrating molecular events that occur between stage 3 and 4, a developmental period required for brown preadipocytes turning into lipid-loaded and thermogenesis active mature BACs. This hypothesis is supported by the following data sets: first, loss of VGLL4 impairs BAC lipogenesis and UCP1 expression; second, the *Vko* BAT gene expression signature was more similar to the differentiated BACs than the undifferentiated brown preadipocytes; third, VGLL4 gain-of-function studies show that activating VGLL4 increased BAT volume and BAC size; fourth, VGLL4 promotes lipogenesis in cultured BACs.

We found that VGLL4 promotes both BAT and WAT lipogenesis *in vivo*. This result differs from studies performed in cultured 3T3L1 cells, which suggested that VGLL4 suppresses adipogenesis ^22^. One possibility is that under physiological conditions, VGLL4 is essential for orchestrating the lipogenesis of adipocytes. However, under harsh chemical adipogenesis conditions, such as the chemical cocktail for inducing 3T3L1 cells differentiation into adipocytes, VGLL4 may be required to block the excessive activity of the adipogenesis factors.

Akt signaling is pivotal for BAT development, because genetically ablation of Akt1 and Akt2 severely diminished BAT growth ^49^. In the neonatal *Vko* BAT, the expression of TEAD1, IGFR and phospho-Akt were all increased, suggestive of VGLL4 impairing BAT development without blunting Akt pathway. PPARγ and CEBPβ are two master regulators of adipogenesis ^13,14,50^. In rodents, CEBPβ-LIP is abundant in fetal BAT and low in adult BAT ^51^, and expressing CEBPβ-LIP in the CEBPβ knockout mice rescued BAT adipogenesis ^35^, implying that CEBPβ-LIP is critical for BAT development. In *Vko* BAT, PPARγ was increased, whereas the expression of two CEBPβ isoforms was differentially regulated, with the larger CEBPβ-LAP isoform remaining unchanged and the smaller CEBPβ-LIP isoform being substantially reduced.

As VGLL4’s partner and one of the terminal effectors of Hippo-YAP pathway, TEAD1 is highly expressed in the P1 BAT and its expression decreases with age. In P0 BAT, TEAD1 bound regions are enriched with CEBP binding motif, suggestive of TEAD1 and CEBP collaborative binding. Using the murine *Casq1* promoter that contains conserved TEAD1 and CEBP binding motifs as a probe, we found that CEBPβ activated *Casq1* expression whereas adding in TEAD1 weakened CEBPβ’s transcriptional activity. This TEAD1 suppression of CEBPβ is likely a direct regulation due to direct protein-protein interactions, because co-IP experiments indicate that these proteins interact with each other. VGLL4 is known to physically interact with TEAD1, and here we show that VGLL4 also interacts with CEBPβ. Although VGLL4 did not affect CEBPβ transcriptional activity, it abolished TEAD1 suppression of CEBPβ. The discovery of VGLL4 alleviating TEAD1 suppression of CEBPβ, together with the aforementioned *Vko* BAT signaling pathways profiling data suggest that VGLL4 and TEAD1 are bridge molecules connecting Hippo-YAP pathway with CEBPβ, one of the core adipogenesis regulators.

YAP/TAZ, TEADs and VGLLs are three categories of crucial transcriptional effectors of the Hippo-YAP pathway. YAP/TAZ interacts with TEADs to regulate target gene expression, and VGLLs function as suppressors of YAP/TEAD1 complex by competing for TEAD ^24^. Published data show that YAP and TAZ suppress BAT lipogenesis, because reducing YAP and TAZ expression in BAT robustly increased BAC lipid content. In the same work, YAP and TAZ were found to serve as mechanosensor proteins in BAT to regulate BAC thermogenesis. Mechanistically, adrenergic stimulation increases cytosolic Ca^2+^that activates actomyosin machinery to cause cellular tension, which in turn increases *Ucp1* expression by activating YAP/TAZ-TEAD complex ^23^. In the current study, our data suggest that VGLL4 has dual roles in the regulation of BAT development: during embryo development, VGLL4 is required for brown preadipocytes differentiating into UCP1 positive brown adipocytes and may regulate adipogenesis by preserving CEBPβ transcriptional activity; from neonate to adulthood, VGLL4 favors lipogenesis and suppresses *Ucp1* expression, and VGLL4 fulfills this function by competing against YAP/TAZ for limited TEAD proteins.

Although UCP1-dependent energy dissipation is critical for BAT thermogenesis ^52^, it is not the only molecular mechanism for BAT to generate heat. UCP1-independent energy dissipation molecular mechanisms, such as Ryr1-mediated Ca^2+^ leak ^53^, also contribute to BAT thermogenesis. Different from YAP/TAZ promoting BAT thermogenesis by increasing UCP1 expression, VGLL4 induction of adult BAT thermogenesis is associated with *Ucp1* down-regulation, thus suggesting that VGLL4 increases BAT thermogenesis by activating UCP1-independent energy dissipation mechanisms. Since *Casq1* encodes a Ca^2+^ buffer protein and it is up-regulated by VGLL4, future studies are required to test whether VGLL4 promotes thermogenesis through activating futile Ca^2+^ recycling ^54^.

In conclusion, our data suggest that Hippo-YAP pathway terminal effector proteins (e.g., TEAD1 and VGLL4) regulate BAT adipogenesis through modulating the transcriptional activity of CEBPβ, thus improving our understanding of how Hippo-YAP pathway regulates BAT development and function.

## Materials and Methods

Refer to the Supplemental Information for Extended Materials and Methods.

### Mice

All animal procedures were approved by the Institutional Animal Care and Use Committee of the Masonic Medical Research Institute. Wild type C57/BL6J mice were used in this study. C57/BL6J mice were originally purchased from Jackson Laboratory and maintained in the AAALAC accredited (#001865) mouse facility of Masonic Medical Research Institute (MMRI). *Vgll4* knockout mouse line ^26^*, Tead1^fb^* knockin ^25^ and BirA mouse lines were published previously ^39^.

### Statistics

Data values (e.g., gene and protein expression data) were expressed as mean ± SD. Normally distributed data were analyzed with student’s t-test (two groups) or one-way ANOVA followed by Tukey’s post hoc test (more than two groups). Non-parametric data (e.g., cell size) were analyzed using Mann-Whitney test (two groups). Prism10 software was used to plot the bar/violin graphs and to perform the statistical analysis.

## Data availability

TEAD1 bioChIP-seq data of postnatal day 0 (P0) mouse brown adipose tissue has been deposited to gene expression omnibus GSE271532. Bulk RNA-seq data of wildtype and Vgll4 knockout P0 mouse brown adipose tissue has been deposited to GSE271533.

## Sources of Funding

**A.** Z. Lin was supported by AHA Scientist Development Grant 15SDG25590001, NIH HL138454-01, and 1R01HL146810. W.T. Pu was supported by NIH R01HL116461 and R01HL146634.

### Author contributions

Z.L. designed and interpreted the study. P.Z., C.W.K. and Z.L. per- formed experiments with the help from A.D., G. W., J.S.K. and S.N.. F. G. analyzed the BAT RNA-seq data. B. D. provided the IBA RNA-seq data and IBA cell line. P. Z., C.W.K. W.T. P and Z. L analyzed the results. Z.L wrote the manuscript. C.W.K. and W.T. P. revised the manuscript.

## Acknowledgment

We thank Jimiao Sun from Boston Children’s Hospital for quantifying the mouse pups lipid content with the magnetic resonance imaging (MRI) data.

