## Supplementary data files for "Vestigial like 4 regulates the adipogenesis of classical brown adipose tissue": Supplementary information_Updated.pdf

Running title: VGLL4 regulates brown adipose tissue development

Pingzhu Zhou[a, *, &], Chase W. Kessinger[b, *], Fei Gu[a], Amanda Davenport[b], Justin S. King[a], Genyu Wang[b], Steven G. Negron[b], Bart Deplancke[c], William T. Pu[a], Zhiqiang Lin[b, #].

a. Boston Children's Hospital, 300 Longwood Ave, Boston, MA, 02115

b. Department of Biomedical Research and Translational Medicine, Masonic Medical Research Institute, 2150 Bleecker St, Utica, NY 13501

c. Swiss Institute of Bioinformatics, CH-1015 Lausanne, Switzerland

&. Current affiliation: Shanghai 411 Hospital, School of Medicine, Shanghai University, 99 Shangda Road, Shanghai, 200444, China.

*P. Z. and C.W.K contributed equally to this work.

**Supplemental information**

**Extended protocol**

*Immortalized brown preadipocyte (IBA) cell line culture and differentiation*

IBA cells were a gift of Dr. Deplancke from Swiss Institute of Bioinformatics. IBAs were cultured in DMEM with 10% FBS and antibiotics, maintained at less than 80% confluence before passaging. Differentiation of IBAs and maintenance of differentiated IBA cells were achieved following a published protocol [1]. To test the effects of VGLL4, adenovirus harboring VGLL4 was added into the IBAs at a dose of 20 multiplicity of infection (MOI) before differentiation. Cells were fixed and stained with Oil Red O six days after differentiation.

*RNA-seq and data analysis*

Total RNA was extracted from the brown adipose tissue of P0 pups using PureLink RNA Mini Kit (Thermo Fisher, 12183020). Polyadenylated RNA from 2 μg total RNA was subsequently purified using Oligo(dT) Dynabeads (Thermo Fisher, 61005). RNA-seq libraries were prepared with ScriptSeq V2 RNA-seq library kit (Illumina, SSV21124). Single-end reads were sequenced using a NextSeq500 sequencer. RNA-seq reads were aligned to mm9 by Tophat [2] and read counts were calculated by Htseq-count [3]. DEseq [4] was next used to perform statistical analysis of differential gene expression. An adjusted P-value of 0.05 was used as cutoff to identify differentially regulated genes.

*Histology and immunostaining*

Brown adipose tissue and inguinal white adipose tissue were fixed in 4% PFA, cryoprotected with 30% sucrose, and embedded in OCT. 10 μm sections were used for Oil Red O staining. H&E staining was performed on paraffin-embedded sections. Immunostaining was performed on cryosections and detected with Alexa-labelled secondary antibodies (Invitrogen). Antibody sources are listed in Supplementary Table 2. EdU was administered intraperitoneally at 200 μg (pregnant dams), 2 hours before tissue collection. EdU was detected with Click-iT chemistry (Invitrogen). Imaging was performed on a Keyence BZ-X800 microscopy system.

*Tead1 bioChIP-seq of interscapular brown adipose tissues*

Tead1 bioChIP-seq was performed as described previously [5]. Briefly, interscapular brown adipose tissues of postnatal day 0 (P0) were harvested. Tissues from 3-4 mice were pooled as one sample, and homogenized with a motor-driven disperser, and then crosslinked in 1% formaldehyde in PBS for 10 minutes at room temperature and subsequently quenched with 125 mM glycine for 5 min. Chromatin was fragmented using a microtip sonicator (Qsonica, S-3000) at 70% amplitude for 8 minutes. Biotinylated Tead1 and bound chromatin was pulled down by incubation with streptavidin beads (ThermoFisher Scientific, 11206D). Bead-bound chromatin was resuspended in SDS elution buffer (50 mM Tris–HCl pH 8,10 mM EDTA, 1% SDS) and incubated overnight at 65 °C to reverse crosslinking. After RNAse A and Proteinase K treatment, ChIP DNA was purified using Qiagen MinElute columns. Libraries were constructed using the KAPA HyperPrep ChIP-seq library preparation kit (Roche, cat# 07962347001). Sequencing (75 nt single end) was performed on an Illumina NextSeq 500.

*Tead1 bioChIP-seq data analysis*

The raw reads from two biological replicate bioChIP-Seq datasets were mapped to the Mus musculus genome (mm9 build) using Bowtie [6] (Ref_Langmead). Alignments with more than one match were removed. MACS2 [7] was used to determine peaks in each replicate with p-value 1e-5 as cutoff. The peak list was further filtered to remove all blacklisted regions. The intersection of peaks from the replicates yielded the reproducible peaks. The intersection and union sets were determined by the Bedtools [8] intersect function. The enriched motifs were determined with Homer de novo motif algorithm analysis [9].

*Luciferase reporter assay*

A 480 bp fragment of mouse *Casq1* genomic DNA was amplified with the following primers that contain KpnI (primer 1) or HindIII (primer 2) restriction sites. Primer 1, 5'-tCAggtaccAAGGACGACACCTTCTGGGCCAGAGG-3'; primer 2, 5'-agcaagcttATGCCCTAGAATCAGAGTCCCACGG-3'. The PCR product was then cloned into pGL2 basic vector and named Casq1_Luci. HEK 293T cells were cultured in 24 well plates for luciferase assay. 100 ng/well indicated plasmids and 10 ng pRLTK internal control vector (Promega) were co-transfected with 1 µL (1 mg/ml) polyethylenimine (PEI). Luciferase activity was measured 24 hours after transfection using the Dual-Luciferase reporter assay system (Promega).

*Magnetic resonance (MR) imaging of total body fat*

Magnetic resonance imaging (MRI) was carried out at Small Animal Imaging Core in

Boston Children's Hospital. Fat volume was analyzed following a published protocol [10].

Acute cold stress

Adult mice were held separately (one mouse/cage) and exposure to an environmental temperature of 4 °C for 24 hours. Core temperate was measured with a RET-3 rectal thermometer probe (world precision Instruments) connected to a TCAT-2DF animal temperature Controller (Physitemp Inc).

In vivo Imaging

Bioluminescence (Luci) imaging was carried out on an IVIS Spectrum instrument with the mice anesthetized with isoflurane. For image capture, mice were placed in the dorsal and ventral imaging positions with image capture 15 minutes post-intraperitoneal injection of D-luciferin (1mg/mouse). For colocalization imaging of the Luci signal from the AAV and microCT segmented BAT, mice were placed in a mouse imaging cassette, which allowed for the coregistration of the Luci signal and microCT datasets[11]. MicroCT imaging was performed on a Quantum GX instrument utilizing a field of view of 60mm with a voxel size of 240µm. For BAT imaging, a voltage of 50 kV and 160 µA was used. MicroCT contrast agent, eXIA 160 (0.1mL/mouse), was injected intravenously four hours before microCT imaging to aid in the identification and segmentation of the interscapular BAT[12]. MicroCT datasets were segmented in ITK-SNAP (v3.8.0), if coregisteration of segmented volumes and Luci signal was needed, it was performed in Living Imaging software (v4.7.3). For higher resolution scans of the interscapular regions, an FOV of 45mm and voxel size of 90 µm was utilized with the same voltage and amperage as above.

AAV9 packaging and administration

AAV9.BCE.GFP, AAV9.BCE.Luciferase, and AAV9.BCE.VGLL4$^{GFP}$ was packaged in 293T cells with AAV9:Rep-Cap and pHelper (pAd deltaF6, Penn Vector Core) and purified and concentrated by gradient centrifugation. AAV titer was determined by quantitative PCR, in which a primer pair amplifying a fragment of the BCE sequence was used. The primers sequences were : 5'-caaggtcaaccctttcctca-3'; 5'-ctgatttggcaccagttcct-3'. 50 µL (1x10^12 GC/ml) AAV was delivered into 3-days-old neonatal pups through subcutaneous injection.

*Adenoviruses*

CMV-GFP adenovirus was described previously [13]. Vgll4-GFP adenovirus was generated by cloning human VGLL4 cDNA with an C-terminal GFP tag into pENTR3C (Invitrogen). These expression cassettes were then transferred to pAd/CMV/V5-DEST using LR clonase (Invitrogen, 11791020).

*Gene expression*

Total RNA was isolated using the Trizol reagent. For quantitative reverse transcription PCR (qRT-PCR), RNA was reverse transcribed (Superscript III) and specific transcripts were measured using SYBR Green chemistry and normalized to either GAPDH or 36B4. Primer sequences are provided in Supplementary Table 1. Western blotting was performed using specific antibodies (see Supplementary Table 2).

**Supplementary Table 1. qRT-PCR primers**

| Gene name | Species | Forward | Reverse |
|---|---|---|---|
| Vgll4 TDU region | Mouse | ACTGCCACCTGTGACCCTGTGGT | GCTCTGGCTCCTTGTAGTTCTTGCCCA |
| Tead1 | Mouse | TACTGCCATCCACAACAAGC | TGCTGCACAAAGGGCTTGAC |
| Tpm1 | Mouse | CGGGCTGAGCTCTCAGAAG | CCGAGTTTCAGCCTCCTTCA |
| Cidea | Mouse | TGCTCTTCTGTATCGCCCAGT | GCCGTGTTAAGGAATCTGCTG |
| Casq1 | Mouse | ATGACCATCCCAGACAAGCC | TTCCTCTGCAAAGGCGACAA |
| Myh9 | Mouse | CCTTGAACGACAACATCGCC | TCCGACAGTACGGAACATGC |
| Smad6 | Mouse | GTGTTGCAACCCCTACCACT | GAATTCACCCGGAGCAGTGA |
| Acsl1 | Mouse | TGCCAGAGCTGATTGACATTC | GGCATACCAGAAGGTGGTGAG |
| Cox7a1 | Mouse | CAGCGTCATGGTCAGTCTGT | AGAAAACCGTGTGGCAGAGA |
| Ucp1 | Mouse | ACTGCCACACCTCCAGTCATT | CTTTGCCTCACTCAGGATTGG |
| Dgat1 | Mouse | GATTGTGGGCCGATTCTTCC | CATACATGAGCACAGCCACC |

| | | | |
|---|---|---|---|
| Fabp4 | Mouse | ACACCGAGATTTCCTTCAAACTG | CCATCTAGGGTTATGATGCTCTTCA |
| Elovl3 | Mouse | TCCGCGTTCTCATGTAGGTCT | GGACCTGATGCAACCCTATGA |
| Prdm16 | Mouse | GCGTGCATCCGCTTGTG | CAGCACGGTGAAGCCATTC |
| Dlk1 | Mouse | CCTGGGTTCTCTGGAAAGGACTG | TGGTTGCGGCTACGATCTCAC |
| AdipQ | Mouse | TGTTCCTCTTAATCCTGCCCA | CCAACCTGCACAAGTTCCCTT |
| Fasn | Mouse | GCTGGCATTCGTGATGGAGTCGT | AGGCCACCAGTGATGATGTAACTCT |
| Plin2 | Mouse | GACCTTGTGTCCTCCGCTTAT | CAACCGCAATTTGTGGCTC |
| Adipor2 | Mouse | CAACTACCAAGGAGATTTGGAGCCCAGC | GCGGGGACATGCCCATAAACCCTTCA |
| 36b4 | Mouse | tgctgaacatctccccttctc | tctccacagacaatgccaggac |
| Cav1 | Mouse | GCGACCCCAAGCATCTCAA | ATGCCGTCGAAACTGTGTGT |
| Cd36 | Mouse | TCCTCTGACATTTGCAGGTCTATC | AAAGGCATTGGCTGGAAGAA |
| Fabp3 | Mouse | agtcactggtgacgctggacg | aggcagcatggtgctgagctg |
| Lpl | Mouse | GGGAGTTTGGCTCCAGAGTTT | TGTGTCTTCAGGGGTCCTTAG |
| Ppargc1b | Mouse | TTGTAGAGTGCCAGGTGCTG | GTGTATCTGGGCCAACGGAA |
| Slc27A1 | Mouse | GCTCAGAACTTCCCAGTCCA | CCACCCACGTACACACAGAA |

**Supplementary Table 2.** **Primary antibodies**

| **Primary antibody** | **Host species** | **Brand (Cat.)** | **Working dilution** |
|---|---|---|---|
| TEAD1 | Mouse | BD biosciences (610923) | 1:1000 for WB; 1:200 for IF |
| PPARγ | Rabbit | CST (#2433) | 1:1000 for WB |
| CEBPβ | Rabbit | CST (#3087) | 1:1000 for WB |
| Akt | Rabbit | CST (#9272s) | 1:1000 for WB |
| Ser473 Phospho Akt | Rabbit | CST (#9271s) | 1:1000 for WB |
| GAPDH | Mouse | Sigma (WH0002597M1) | 1:1000 for WB |
| UCP1 | Rabbit | Abcam (ab10983) | 1:1000 for WB; 1:200 for IF |
| IGF1R | Chicken | Novus Biologicals | 1:1000 for WB |
| ATGL | Rabbit | CST (2138S) | 1:1000 for WB |
| GFP | Goat | Rockland (600-101-215) | 1:1000 for WB |
| Flag | Mouse | Sigma (F1804) | 1:1000 for WB |
| HA | Rabbit | CST (C29F4) | 1:1000 for WB |

#### Supplemental Figures

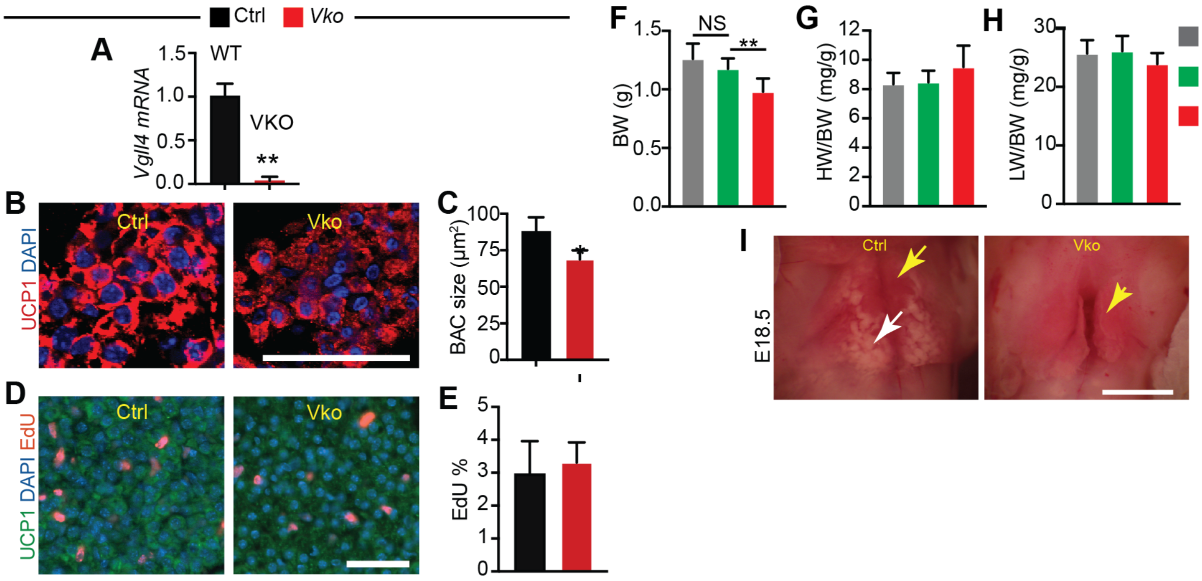


**Supplementary Figure 1. Related to Figure 1.**

**A.** BAT Vgll4 mRNA expression. E18.5 interscapular BAT was collected for mRNA measurement by qRT-PCR. TDU1 domain of *Vgll4* was not detectable in the Vgll4 mutant transcript. **B**. Immunofluorescence images of the BAT sections. **C**. Measurement of P0 brown adipocytes (BACs) size. Student t-test, *P<0.05. For each mouse, 50 BACs were measured, and the average BAC size was presented. For each group, n=4. **D-E**. Measurement of BAT cell proliferation rate using BAT tissues from E18.5 mouse embryo. H, representative images of EdU and UCP1 immunofluorescence staining. I, percentage of UCP1+ cells that are EdU+. N=3. F and H, scale bar=50 μm. **F**. Body weight at E18.5. **G**. Heart to body weight ratio at E18.5. **H**. Lung to body weight ratio. A, B and C, n=5 for each group. **I**. Gross morphology of E18.5 mouse embryo BAT. Bar=2 mm. Yellow arrow indicates BAT; White arrow indicates white adipose tissue covering BAT.

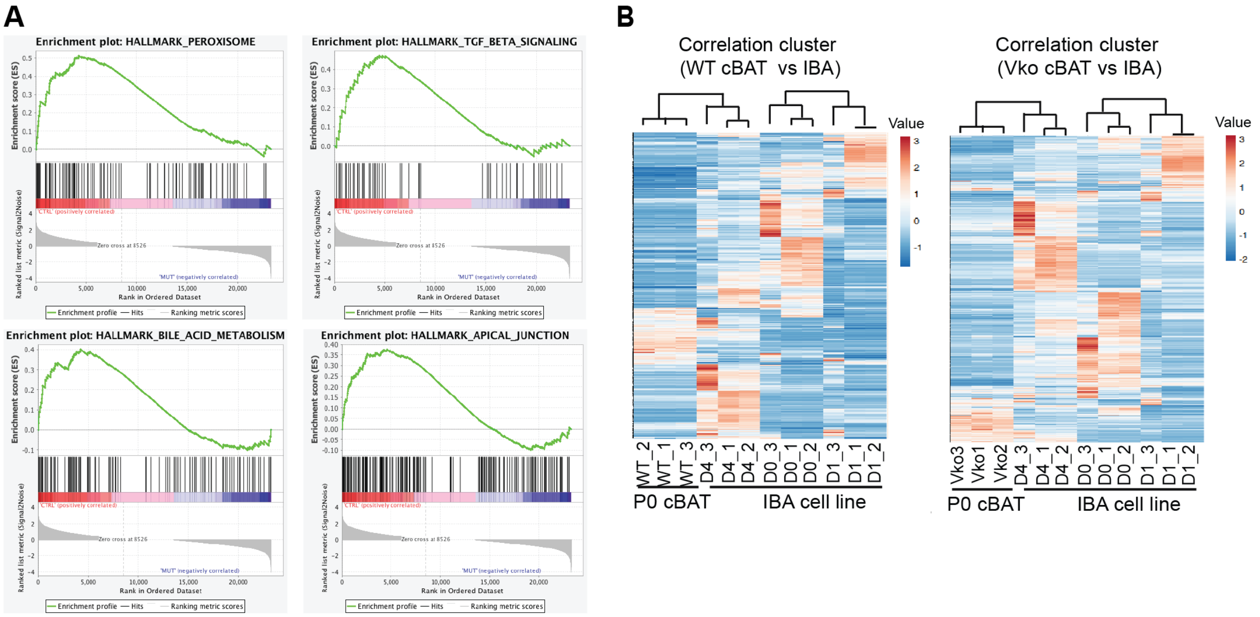


| NAME | SIZE | NOM p-val | FDR q-val |
|---|---|---|---|
| HALLMARK_MYOGENESIS | 196 | 0 | 0 |
| HALLMARK_ADIPOGENESIS | 191 | 0 | 0 |
| HALLMARK_OXIDATIVE_PHOSPHORYLATION | 176 | 0 | 0.00017 |
| HALLMARK_FATTY_ACID_METABOLISM | 147 | 0 | 0.00018 |
| HALLMARK_PEROXISOME | 96 | 0 | 0.0002 |
| HALLMARK_TGF_BETA_SIGNALING | 53 | 0.00562852 | 0.01664096 |
| HALLMARK_BILE_ACID_METABOLISM | 109 | 0.00371058 | 0.03022808 |
| HALLMARK_APICAL_JUNCTION | 188 | 0 | 0.02821333 |

**Supplementary Figure 2. Related to Figure 2.**

**A.** Gene set enrichment analysis (GESA). GESA was carried out with the Deseq2-derived gene expression data set. The top eight gene sets enriched in wild-type cBAT were listed. **B**. Gene expression comparison between P0 BAT and differentiated brown adipocytes at different stages. Correlation cluster analysis was carried out using BAT and published brown adipocytes (IBA cells) RNA seq data. Brown adipocytes differentiation stages: Day 0 (D0), Day 1 (D1) and Day 4 (D4). Unit variance scaling value was applied to rows. Both rows and columns are clustered using correlation distance and average linkage.

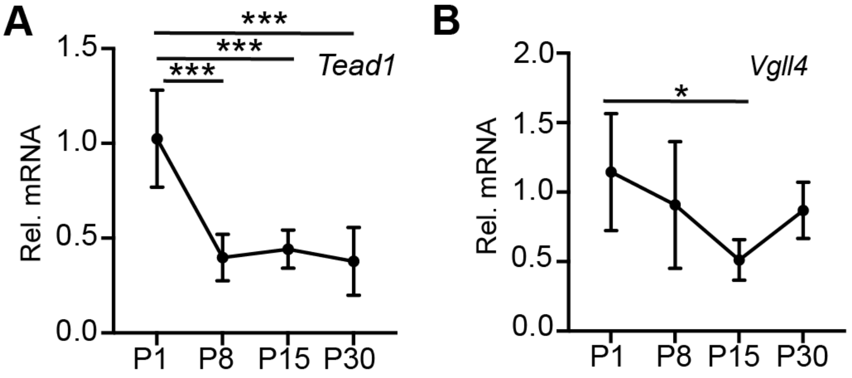


**Supplementary Figure 3. Related to Figure 3.**

Measurement of *Tead1* (**A**) and *Vgll4* (**B**) mRNA levels. Interscapular BAT collected from mice of different ages was used for qRT-PCR measurement. All the tested mRNAs were normalized by endogenous 36B4 mRNA. One-way ANOVA with post-hoc Tukey Honestly Significant Difference test, *, $p<0.01$; ***, $p<0.001$. For each group, n=5-6.

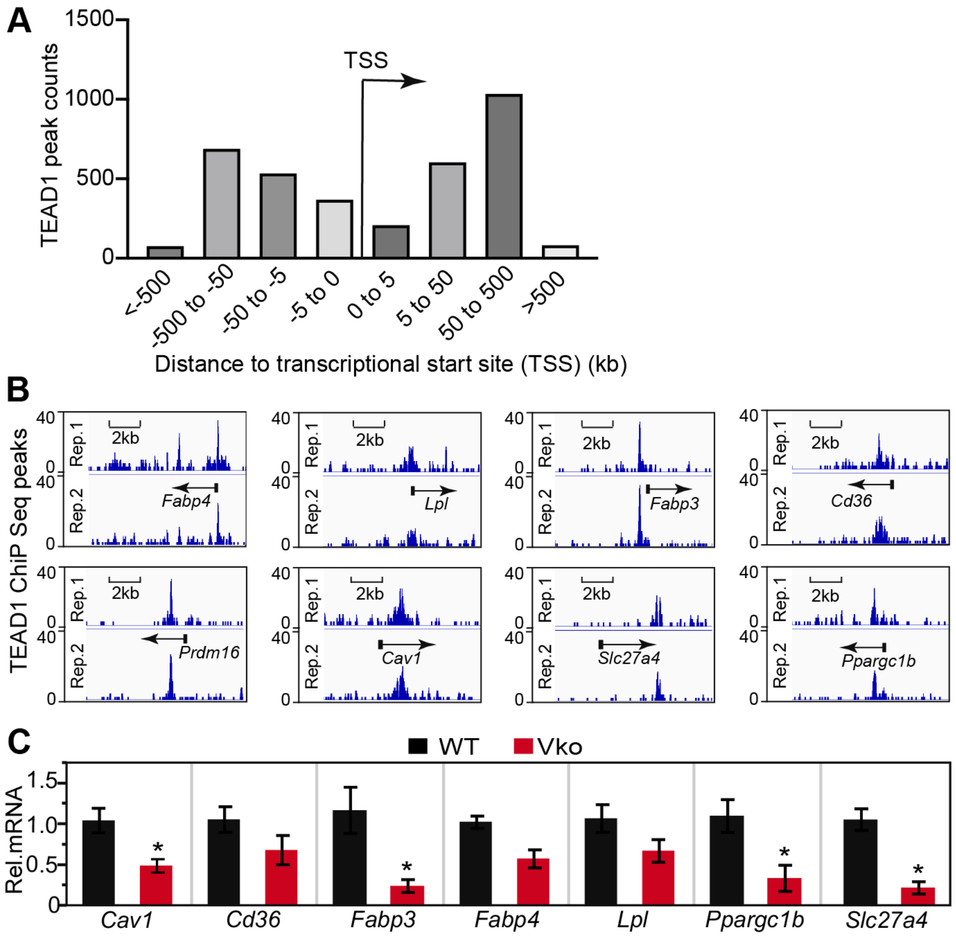


**Supplementary Figure 4. Related to Figure 4.**

**A.** Distribution of TEAD1 bound regions relative to the nearest TSS across the mouse genome (x-axis, number of genes bound by TEAD1; y-axis, distance relative to the TSS from –500 kb to +500 kb) **B.** Genome browser view showing ChIP-seq of TEAD1 occupancy near the transcriptional start site (TSS) or gene body of prominent adipogenesis genes. ChIP-seq data from two biological replicates are shown. **C.** qRT-PCR measurement of gene expression. P0 control and *Vko* BAT mRNA was used for qRT-PCR analysis. All the tested mRNAs were normalized by endogenous 36B4 mRNA. N=4. Student t-test, *, P<0.05.

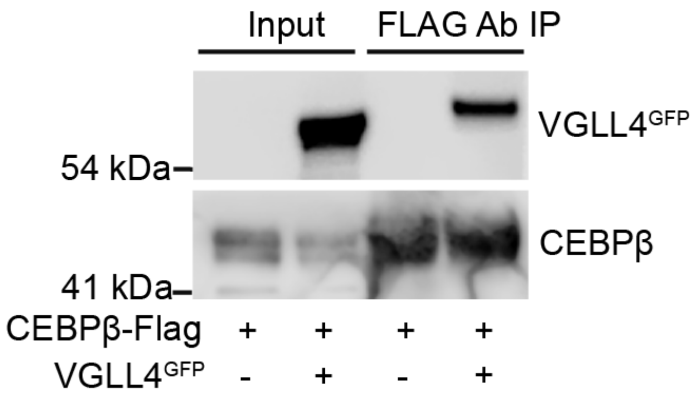


**Supplementary Figure 5. Related to Figure 5.** CEBPβ Co-immunoprecipitation (Co-IP) assay. Indicated plasmids were transfected into HEK293T cells. Flag antibody was used for Co-IP.

**A**

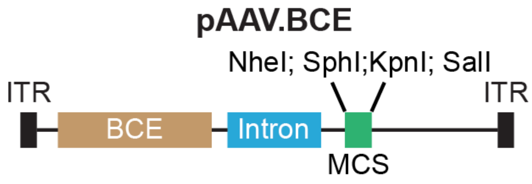


**B**

**BCE sequence**

5’-gcatgccaatttatagtgccgtcactaacagtactgatactttaacatgctaagttta-
aagtgtgtgctatattaattgtaagattggtgaagagaggtgttatcagatggaagctgc
acatttctggattaatgtggttaaatgtatcttctcctgtgattactgtctttatttcttcttttaaa
atattgtcatttggacatctatctgtatagctacgccctgacacgtcctcctggagacaga
taagaagttacgacgggaggagcagatggaggcaaagcgctgtgatgcttttgtggtt
tgagtgcacacatttgttcagtgattctgtgaaatgagtgagcaaatggtgaccgggtgc
cctgtaaatggtgttctacatcttaagagaagaacacggacactaggtaagtgaagctt
gctgtcactcctctacagcgtcacagagggtcagtcacccttgaccacactgaactagt
cgtcacctttccactcttcctgccagaagagcagaaatcagactctctggggatatcag
cctcacccctactgctctctccattatgaggcaaactttctttcacttcccagaggctctgg
gggcagcaaggtcaaccctttcctcagactctagtctcggaggagatcagatcgcgct
tattcaagggaaccagcccctgctctgcgccctggtccaaggctgttgaagagtgaca
aaaggcaccacgctgcggggacgcgggtgaagcccctctgtgtgtcctctgggcata
atcaggaactggtgccaaatcagaggtgatgtggccagggctttgggagtgacgcgc
ggctgggaggcttgcgcacccaaggcacgcccctgccaagtcccactagcagctctt
tggagacctgggccggctcagccacttcccccagtccctcctccggcaaggggctat
atagatctcccaggtcagggcgcag-3’

**Supplementary Figure 6. Related to Figure 6.**

**A**. Schematic view of pAAV.BCE construct. ITR, inverted terminal repeat sequence. MCS, multiple cloning site. Unique restriction enzyme cutting sites were listed. BCE, brown adipocyte cis-regulatory element. **B.** BCE sequence. Black letters indicate UCP1 enhancer, brown letters indicate UCP1 mini promoter.

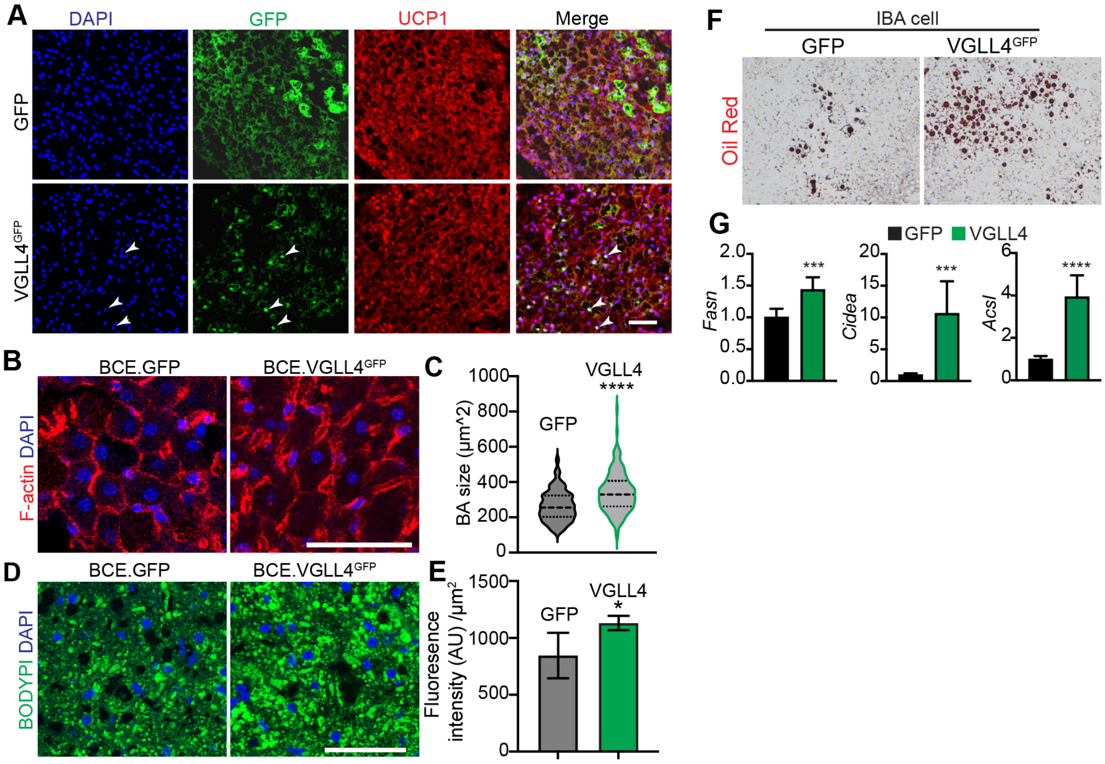


**Supplementary Figure 7. Related to Figure 7.**

**A.** Immunofluorescence images of BAT**.** BAT sections were co-stained with UCP1 and GFP antibodies**.** White arrowheads indicate nuclei-localized VGLL4. Scale bar = 50 μm. **B**. Fluorescence images of BAT. BAT sections were stained with phalloidin (F-actin) and DAPI. Scale bar = 50 μm. **C**. Quantification of P42 brown adipocytes size. Mann-Whitney test, ****, p<0.05. For each group, 150 cells from 3 animals were measured. **D.** BODIPY stained BAT sections. Scale bar = 50 μm. **E.** Quantification of BODIPY fluorescence intensity. BODIPY fluorescence intensity was normalized to tissue area. AU, arbitrary unit. BCE.GFP, n=4; BCE.VGLL4, n=3. **F**. Representative Oil Red O staining of IBA cells treated with brown adipocyte differentiation cocktail chemicals. **G**. qRT-PCR measurement of lipogenesis genes. All the tested mRNAs were normalized by endogenous 36B4 mRNA. N=5. Student t-test, ***, P<0.001.

### Original western blots

**Figure 1B**

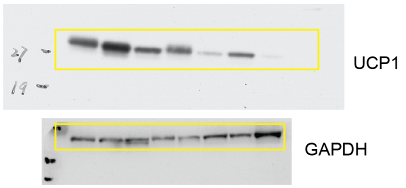


**Figure 3A**

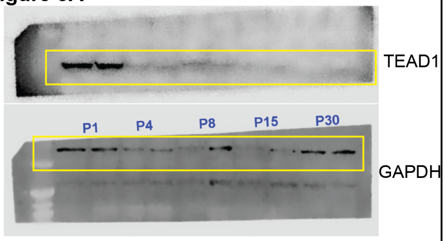


**Figure 3E**

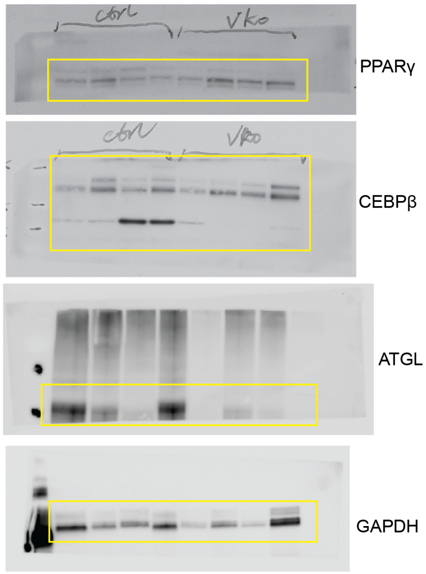


**Figure 3C**

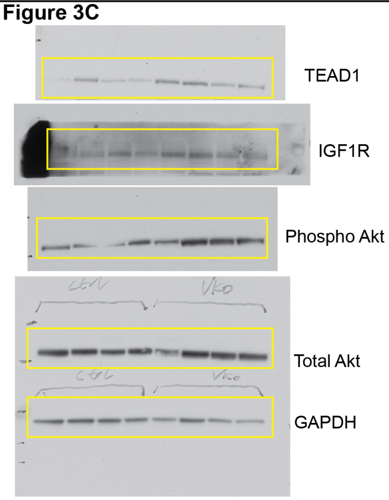


**Figure 5F**

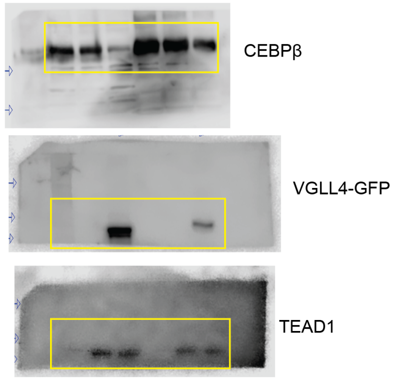


**Figure 7E**

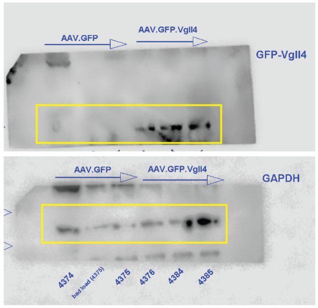


**Supplementary Figure 5**

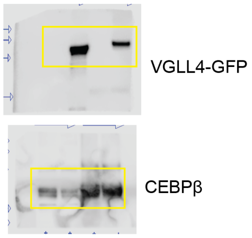
